# The genetic basis of energy conservation in the sulfate-reducing bacterium Desulfovibrio alaskensis G20

**DOI:** 10.1101/005694

**Authors:** Morgan N. Price, Jayashree Ray, Kelly M. Wetmore, Jennifer V. Kuehl, Stefan Bauer, Adam M. Deutschbauer, Adam P. Arkin

## Abstract

Sulfate-reducing bacteria play major roles in the global carbon and sulfur cycles, but it remains unclear how reducing sulfate yields energy. To determine the genetic basis of energy conservation, we measured the fitness of thousands of pooled mutants of *Desulfovibrio alaskensis* G20 during growth in 12 different combinations of electron donors and acceptors. We show that ion pumping by the ferredoxin:NADH oxidoreductase Rnf is required whenever substrate-level phosphorylation is not possible. The uncharacterized complex Hdr/flox-1 (Dde_1207:13) is sometimes important alongside Rnf and may perform an electron bifurcation to generate more reduced ferredoxin from NADH to allow further ion pumping. Similarly, during the oxidation of malate or fumarate, the electron-bifurcating transhydrogenase NfnAB-2 (Dde_1250:1) is important and may generate reduced ferredoxin to allow additional ion pumping by Rnf. During formate oxidation, the periplasmic [NiFeSe] hydrogenase HysAB is required, which suggests that hydrogen forms in the periplasm, diffuses to the cytoplasm, and is used to reduce ferredoxin, thus providing a substrate for Rnf. During hydrogen utilization, the transmembrane electron transport complex Tmc is important and may move electrons from the periplasm into the cytoplasmic sulfite reduction pathway. Finally, mutants of many other putative electron carriers have no clear phenotype, which suggests that they are not important under our growth conditions.

## Introduction

Sulfate-reducing bacteria are major players in the remineralization of fixed carbon and in the global sulfur cycle, but their energy metabolism remains poorly understood. Research on the mechanism of sulfate reduction has focused on members of the genus Desulfovibrio, which are relatively easy to culture in the laboratory. Sulfate reduction is best studied in the strain *Desulfovibrio vulgaris* Hildenborough (Keller and Wall, 2011), but the Desulfovibrio genus is quite diverse. We are studying the energy metabolism of *Desulfovibrio alaskensis* G20 (formerly *D. desulfuricans* G20), for which a large collection of mutants is available (Kuehl et al., 2014). G20 is a derivative of the G100A strain that was isolated from an oil well in Ventura County, California (Wall et al., 1993). |Only 1,871 of 3,258 proteins in the genome of *D. alaskensis* G20 (Hauser et al., 2011) have orthologs in *D. vulgaris* Hildenborough.

The key mystery of sulfate reduction is: how does it lead to net ATP production? Sulfate must be activated to adenosine 5’-phosphosulfate (APS), which costs two ATP (Figure 1A). Desulfovibrios can oxidize lactate to pyruvate and then to acetyl-CoA, which is then converted to acetate while converting one ADP to one ATP (Figure 1B). Reducing sulfate to sulfide requires 8 electrons, while oxidizing lactate to acetate yields 4 electrons, so lactate and sulfate are utilized at a molar ratio of 2:1. Thus, the ATP from substrate-level phosphorylation is balanced out by the cost of activating sulfate (Peck, 1960).

**Figure 1.**
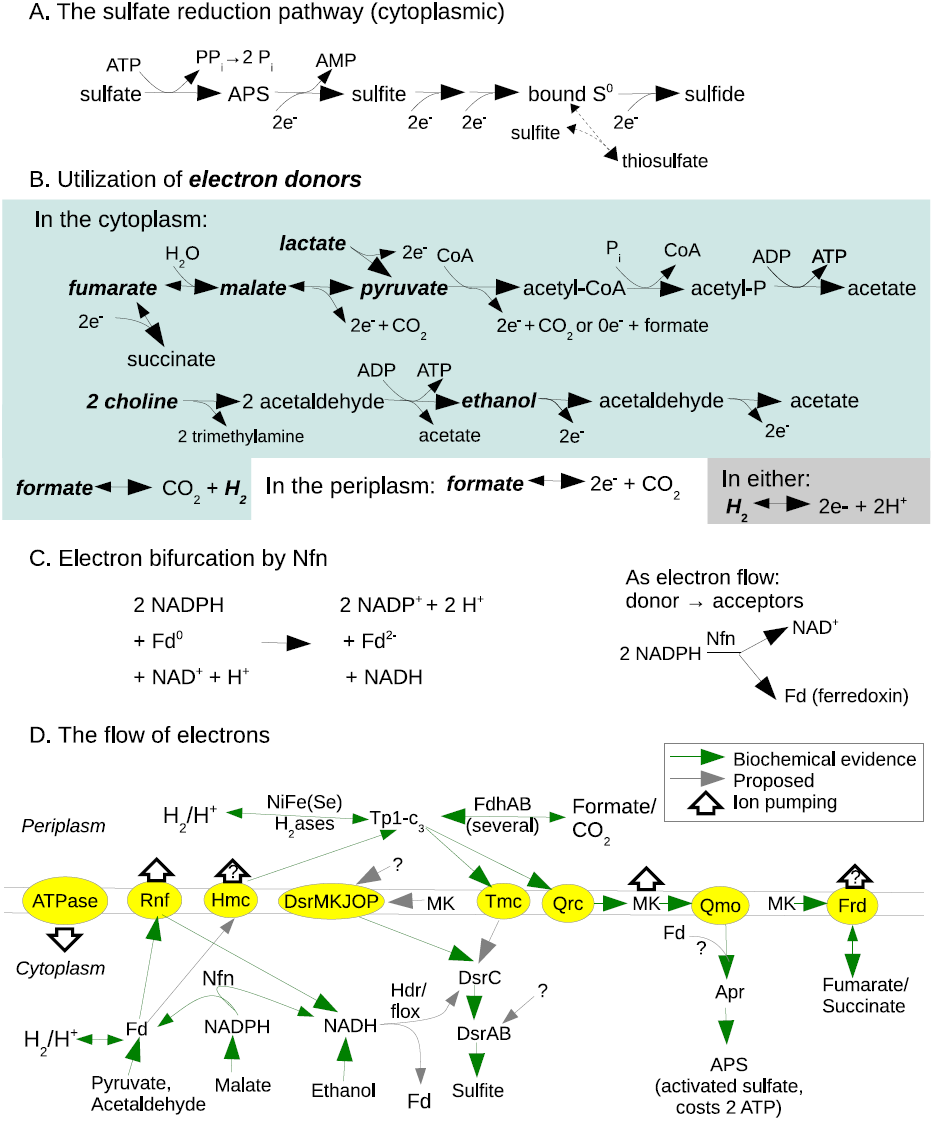
Overview of energy metabolism of *D. alaskensis* G20, including (A) sulfate reduction, (B) utilization of electron donors (which are in bold), (C) illustration of electron bifurcation by Nfn, and (D) overview of electron flow. Fd is ferredoxin; MK is menaquinone; Tp1-c_3_ is type 1 cytochrome c_3_; Apr is APS reductase.

Furthermore, *D. alaskensis* G20 can grow via sulfate reduction while oxidizing ethanol, formate, or molecular hydrogen, and oxidation of these substrates is not expected to lead to any substrate-level phosphorylation (Figure 1B). In these conditions, reducing a molecule of sulfate gives a loss of 2 ATP, which must be made up for by ATP synthase, which can convert an ion gradient into chemical energy. It is estimated that another strain, D. *vulgaris* Marburg, obtains about 1 net mole of ATP per mole of sulfate converted to sulfide while oxidizing hydrogen (Badziong and Thauer, 1978). The stoichiometry of the ATP synthase reaction varies across bacteria and even across growth conditions, with a typical range of 2–4 protons translocated per ATP formed (Tomashek and Brusilow, 2000). If, in Desulfovibrios, ATP synthase moves 3 protons per ATP, then to yield 1 net ATP, around 9 protons must be pumped per 8 electrons used for reducing sulfate, or roughly 1 proton pumped per electron.

There are many theories for how the proton gradient might be formed. Key redox complexes such as the pyruvate:ferredoxin oxidoreductase which oxidizes pyruvate to acetyl-CoA, APS reductase, and sulfite reductase are located in the cytoplasm and are not associated with a membrane, which would seem to preclude proton pumping by a membrane-bound electron transport chain. This, together with the tendency of Desulfovibrios to produce a “burst” of hydrogen at the beginning of batch growth on lactate/sulfate media, led to the hydrogen cycling model (Odom and Peck, 1981). During hydrogen cycling, electrons would move from the electron donor to cytoplasmic ferredoxin, which is believed to be a major cytoplasmic electron carrier, to a cytoplasmic hydrogenase, which combines two electrons with two protons to evolve H_2_. The hydrogen then diffuses to the periplasm, where a periplasmic hydrogenase oxidizes it to produce two periplasmic protons and two electrons. The electrons would then move through transmembrane complexes (there are many candidates in the *Desulfovibrio* genomes) into the cytoplasm to reduce sulfate. In principle this mechanism can pump 2 protons per molecule of H_2_, or 1 proton per electron transferred from ferredoxin, or 8 protons per molecule of sulfate. However, genetic evidence suggests that hydrogen cycling is not required for sulfate reduction by Desulfovibrios. For example, in *D. alaskensis* G20, mutants of type 1 cytochrome c_3_ (Tp1-c_3_, also known as *cycA*), which is the major periplasmic electron carrier, grow in lactate/sulfate media but cannot utilize hydrogen as an electron donor (Rapp-Giles et al., 2000; Li et al., 2009; Keller et al., 2014). Similarly, in *D. vulgaris* Hildenborough or *D. gigas*, mutants of *cycA* or of various hydrogenases grow in lactate/sulfate media (Sim et al., 2013; Morais-Silva et al., 2013). Thus, uptake of hydrogen in the periplasm is not required to obtain energy by sulfate reduction when oxidizing lactate.

Another potential mechanism for forming a proton gradient is formate cycling. In *D. vulgaris* Hildenborough, formate dehydrogenases are present only in the periplasm, but formate could be formed in the cytoplasm by pyruvate-formate lyase, which generates acetyl-CoA and formate from pyruvate and coenzyme A. The genome of *D. alaskensis* G20 encodes these enzymes and also a putative cytoplasmic formate:hydrogen lyase that may convert cytoplasmic formic acid to H_2_ and CO_2_ or vice versa (Pereira et al., 2011). In either case, formic acid could diffuse through the cytoplasmic membrane and be reoxidized in the periplasm. As with hydrogen cycling, formate cycling would pump one proton per electron transferred. There is evidence that formate cycling contributes to energy production in *D. vulgaris* Hildenborough, as knockouts of formate dehydrogenases had reduced growth in lactate/sulfate media (da Silva et al., 2013). In D. *alaskensis* G20, during growth on lactate/sulfate media, a *cycA* mutant accumulated formate, but the parent strain did not (Li et al., 2009), which suggests that formate might normally be formed and then immediately reoxidized.

In all of these cycling models, the electrons return to the cytoplasm to reduce sulfate via a transmembrane electron transfer protein. The genomes of both *D. vulgaris* Hildenborough and *D. alaskensis* G20 contain a variety transmembrane redox complexes that could return electrons to the cytoplasm (Pereira et al., 2011). In particular, the Qrc complex can transfer electrons from the periplasmic Tp1-c_3_ to menaquinone, an electron carrier in the membrane, and the Qmo complex is believed to transfer electrons from menaquinol to APS reductase, which then reduces APS to sulfite (Venceslau et al., 2010; Ramos et al., 2012; Krumholz et al., 2013). Furthermore, because the reduction and oxidation of menaquinone may involve adding protons from the cytoplasm and removing protons into the periplasm, the combination of Qrc and Qmo could create a proton gradient. Qmo is essential for sulfate reduction (Zane et al., 2010), but Qrc (previously known as mopB) is primarily needed for hydrogen or formate oxidation (Li et al., 2009; Keller et al., 2014). A path from the periplasm to sulfite reduction is less clear, but the transmembrane complex DsrMKJOP interacts with DsrC and hence is suspected to send electrons from the periplasm and/or from menaquinol to DsrC (Grein et al., 2010; Pereira et al., 2011). (A potential issue with this model is that DsrMKJOP appears not to accept electrons from periplasmic hydrogenases or Tp1-c_3_ (Pires et al., 2006).) The DsrC protein is part of the dissimilatory sulfite reductase (DsrABC) but is also believed to disassociate from DsrAB and act as a diffusible electron carrier for two of the six electrons that are required to reduce sulfite to sulfide (Oliveira et al., 2008). So, DsrMKJOP in combination with DsrC and DsrAB could use electrons from the periplasm to reduce sulfite to sulfide. *D. vulgaris* Hildenborough and *D. alaskensis* G20 also contain transmembrane redox complexes Hmc (high-molecular weight cytochrome, with a 16-heme periplasmic subunit) and Tmc (with a periplasmic type II cytochrome c_3_ subunit), which are believed to accept electrons from Tp1-c_3_ and transfer them across the membrane (Pereira et al., 1998, 2006; Quintas et al., 2013).

There are also a variety of alternatives to the cycling models. The genomes of both *D. vulgaris* Hildenborough and *D. alaskensis* G20 encode Rnf, an ion-pumping ferredoxin:NADH oxidoreductase, which can generate an ion gradient without moving electrons to the periplasm (Biegel et al., 2011). Although the best-studied Rnf complexes pump sodium ions, Rnf from *Clostridum ljungdahlii* appears to pump protons (Tremblay et al., 2013). As the Desulfovibrio Rnf is distantly related to all characterized Rnf, the ion pumped by Rnf in Desulfovibrios cannot be guessed. Here we will show that in *D. alaskensis* G20, Rnf is important for growth under sulfate-reducing conditions with a variety of electron donors.

Additional possibilities for energy conservation arise because the roles of DsrMKJOP, Hmc, and Tmc are not fully understood. They might be able to move electrons between the cytoplasm and menaquinone without involving Tp1-c_3_. The exchange of protons between menaquinol, the cytoplasm, and the periplasm could also create a proton gradient.

Yet another potential mechanism of energy conservation arises from electron bifurcation, which is the transfer of electrons from a single source to two different sources. For example, the genomes of *D. vulgaris* Hildenborough and *D. alaskensis* G20 both encode homologs of the electron-bifurcating transhydrogenase Nfn, which couples electron transfer from NADPH to NAD+, which is energetically favorable, to electron transfer from NADPH to ferredoxin, which is energetically unfavorable, i.e., 2 NADPH + Fd^0^ + NAD^+^ ↔ 2 NADP^+^ + Fd^2–^ + NADH + H^+^ (Wang et al., 2010) (Figure 1C). Both genomes also encode the putative redox complex Hdr/flox, which is proposed to bifurcate electrons from NADH to a heterodisulfide such as DsrC (favorable) and to ferredoxin (unfavorable) (Pereira et al., 2011). Based on our data, we propose that these electron bifurcations allow an increased yield of reduced ferredoxin, which can be used by Rnf to pump additional ions into the periplasm. The combination of Hdr/flox and Rnf was previously proposed to be involved in energy production during pyruvate fermentation by *D. alaskensis* G20 (Meyer et al., 2014).

Electron bifurcation might be involved more directly in sulfate reduction: the QmoB subunit of the Qmo complex, which is essential for sulfate reduction, is homologous to HdrA, a subunit of heterosulfide reductases from methanogens that are believed to perform electron bifurcations (Ramos et al., 2012). Although Qmo interacts with APS reductase in vitro, electron transfer from menaquinol analogs to APS reductase was not reconstituted, so it is proposed that a second electron donor might be required (Ramos et al., 2012). For example, Qmo might reduce APS reductase while oxidizing both menaquinol and a more energetically favorable electron donor such as ferredoxin. This would be an electron bifurcation in reverse (a confurcation).

Overall, the genomes of Desulfovibrios reveal a wide variety of potential mechanisms by which energy could be conserved, and it remains unclear which of them are important for generating energy. To address this issue, we measured the growth of thousands of pooled mutants of *D. alaskensis* G20 with 12 combinations of electron donors and electron acceptors. We also verified the phenotypes of key redox complexes by growing mutants individually. We found that Rnf, Nfn, and Hdr/flox are involved in energy conservation in some sulfate-reducing conditions. We believe that this is the first evidence of a role for these complexes in energy conservation during sulfate reduction. We found that formate utilization requires the formation of H_2_; we propose that this is necessary to allow the reduction of ferredoxin. We found that mutants in Tmc are deficient in hydrogen oxidation, which is consistent with a biochemical study (Pereira et al., 2006). Finally, mutants in many putative electron carriers lacked clear phenotypes, which suggests that they are not important for energy conservation (although we cannot rule out genetic redundancy). In particular, we found no evidence of energy conservation by molecular cycling. Based on our genetic data, we propose an overview of electron flow and energy conservation in *D. alaskensis* G20 (Figure 1D).

## Results and Discussion

### Genome-wide fitness data

We used a collection of transposon mutants of *D. alaskensis* G20 that have been mapped and tagged with DNA barcodes (Kuehl et al., 2014). The DNA barcodes allow us to measure the relative fitness of strains with mutations in most of the non-essential genes in the genome during pooled (competitive) growth. Specifically, for each energetic condition, we grew two different pools of mutants separately and we used the strains’ barcodes to measure how the abundance of each strain changed during growth in that condition. The fitness of a strain is defined as the change of abundance on a log_2_ scale, i.e., log_2_(end / start). The data is normalized so that strains that grow better than the typical strain have positive fitness and strains that grow poorly have negative fitness (Price et al., 2013). The fitness of a gene is defined as the average of the fitness values for the relevant strains.

The fitness data includes 6,500 strains that have insertions within 2,369 of the 3,258 protein-coding genes. We lack data for genes that are required for sulfate reduction or lactate oxidation. This is expected because most of the mutants were isolated on lactate/sulfate media (Kuehl et al., 2014). Otherwise, if we lack fitness data for one gene of interest, we usually have fitness data for functionally-related genes in the same operon, but there are a few exceptions. The energy-related genes that we lack fitness data for are described in Text S1.

We assayed the growth of pools of mutants in 12 different combinations of electron donors and electron acceptors, including growth with sulfate as the electron acceptor and 8 different electron donors (pyruvate, choline, lactate, fumarate, malate, ethanol, hydrogen, or formate). We also studied growth with sulfite as the electron acceptor and pyruvate or lactate as electron donors; with thiosulfate as the electron acceptor and lactate as the electron donor; and pyruvate fermentation. This includes a total of 49 genome-wide fitness experiments. Some conditions were repeated with or without yeast extract or vitamins added, with a different reductant, or with a different buffer (Dataset S1). For experiments with hydrogen or formate as the electron donor, acetate was added to the media to serve as the carbon source. In the typical experiment, the mutant pools doubled 4.7 times (growth from OD_600_ = 0.02 to 0.6). A defined ethanol/sulfate medium gave the lowest growth yield (2.5–3.1 doublings), while a pyruvate/sulfate medium with yeast extract gave the best growth yield (5.6 doublings).

We observed strong phenotypes in a subset of energetic conditions for many energy-related genes, including genes that are expected to be involved in the utilization of specific electron donors or acceptors and genes encoding many electron transfer complexes. As can be seen in Figures 2–4, phenotypes for genes in the same complex were consistent, and similar energetic conditions usually gave similar fitness values. We will first discuss the pathways for the utilization of electron donors, then the pathways for utilization of electron acceptors, and finally we will discuss the role of electron transport complexes in energy conservation.

**Figure 2.**
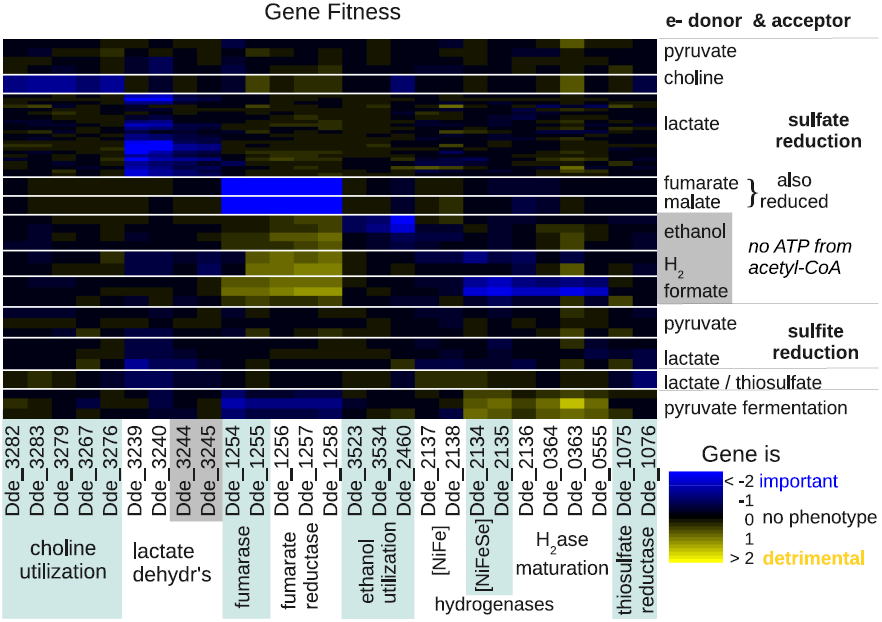
Heatmap of fitness data for genes relating to the utilization of various electron donors or acceptors, across 12 energetic conditions.

**Figure 3.**
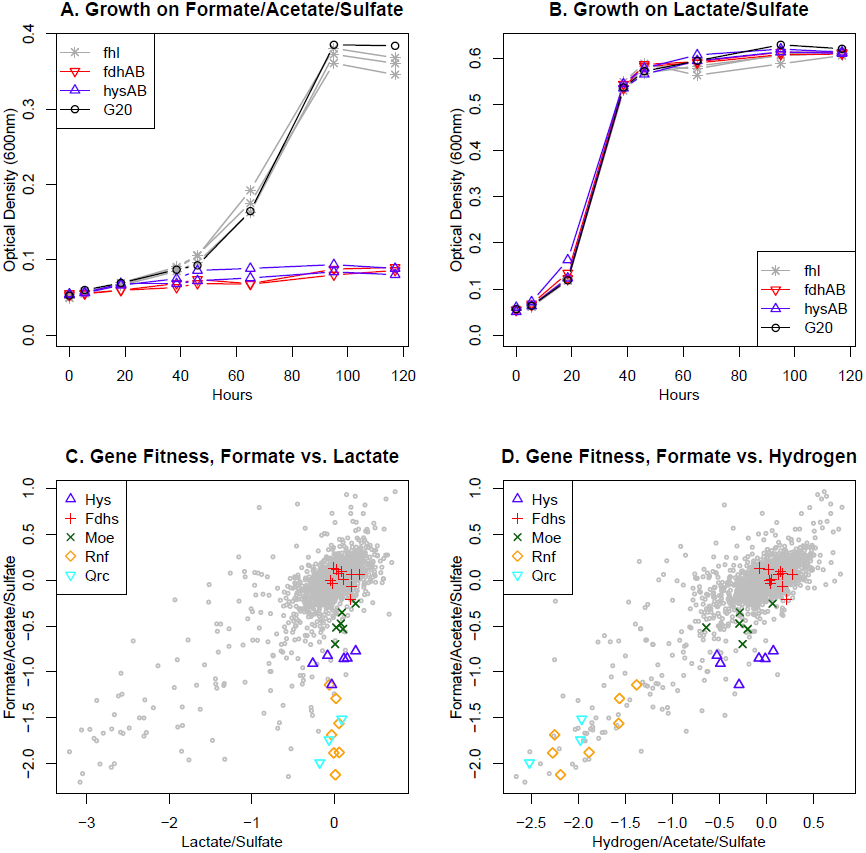
Evidence for hydrogen production during growth on formate/acetate/sulfate medium. **(A & B)** Growth of mutants in the periplasmic [NiFeSe] hydrogenase (*hysAB*), a periplasmic formate dehydrogenase (*fdhAB*, Dde_0717:Dde_0718), formate:hydrogen lyase (*fhl*), or of the parent strain. Growth was measured for 2–3 different mutants in each complex, and each point shows the average across three cultures for a strain. **(A)** Growth in 50 mM formate, 10 mM acetate, and 30 mM sulfate. **(B)** Growth in 60 mM lactate and 30 mM sulfate. **(C & D)** Comparisons of gene fitness with formate, lactate, or hydrogen as electron donor. “Fdhs” includes periplasmic formate dehydrogenases and formate:hydrogen lyase. “Moe” includes molybdopterin synthesis genes (Dde_0709, Dde_1390, Dde_0249, Dde_2352, Dde_0230, Dde_3228). “Hys” includes *hysAB* and maturation genes (Dde_2136, Dde_0364, Dde_0363, Dde_0555). The fitness data is the average of two independent experiments for each of two pools of mutants. Both growth and fitness experiments were performed in MO media with 1 mM sulfide (as reductant) and no added yeast extract or vitamins.

**Figure 4.**
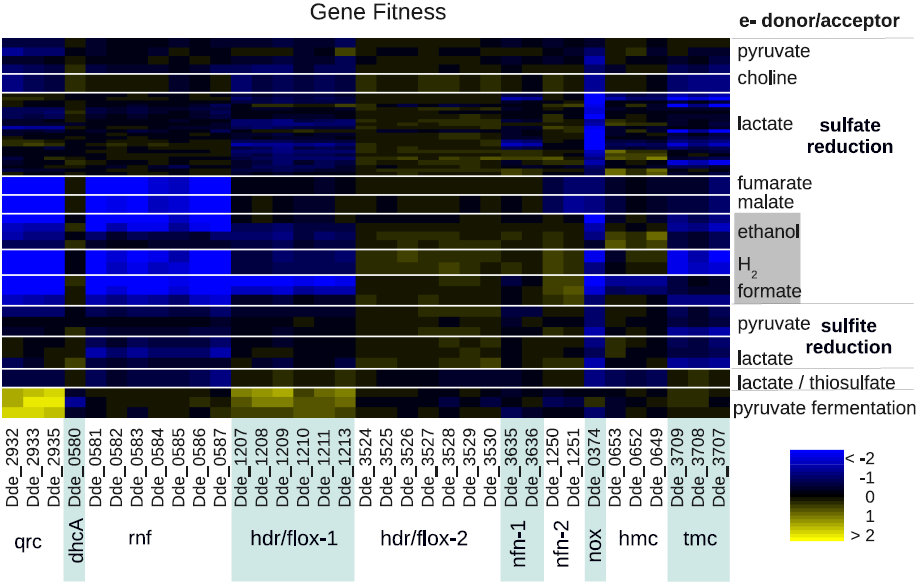
Heatmap of fitness data for central electron transport complexes across 12 energetic conditions.

## Utilization of electron donors

*Pyruvate.* As discussed above, we lack data for many of the genes that are required for lactate utilization, and as lactate is oxidized to pyruvate, this may also explain why we did not identify genes that were specifically important for pyruvate utilization. In particular, pyruvate is expected to be oxidized by pyruvate:ferredoxin oxidoreductase (Dde_3237), which we lack data for. Pyruvate could also be converted to acetyl-CoA and formate by pyruvate:formate lyase (Dde_3039, Dde_3055, or Dde_1273). Dde_1273 and its putative activating enzyme Dde_1272 had a moderate fitness defect in some defined lactate/sulfate experiments (mean fitness = −0.7 to −1.1), but were not important during growth on pyruvate (fitness = −0.2 to 0). The other pyruvate-formate lyases lacked phenotypes (Figure S1). Dde_1273 is related to choline:trimethylamine lyase and glycerol dehydratase (Craciun and Balskus, 2012; Raynaud et al., 2003), so given its phenotypes, we suspect that Dde_1273 is not a pyruvate:formate lyase. Overall, pyruvate is probably consumed primarily by pryuvate:ferredoxin oxidoreduct ase.

**Figure S1.**
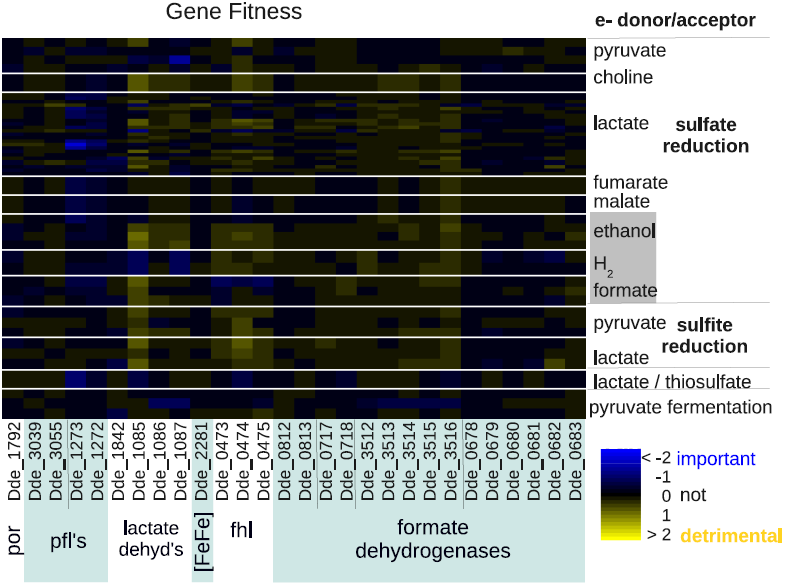
Heatmap of fitness data for genes that might relate to the utilization of various electron donors, but which lack strong phenotypes across 12 energetic conditions. por - pyruvate:ferredoxin oxidoreductase. pfl - pyruvate:formate lyase. [FeFe] - periplasmic iron only hydrogenase. fhl - formate:hydrogen lyase.

*Choline.* Choline oxidation occurs in a microcompartment that is encoded by a large gene cluster (Craciun and Balskus, 2012; Kuehl et al., 2014). This cluster includes choline:trimethylamine lyase Dde_3282 (Craciun and Balskus, 2012), which splits choline to trimethylamine and acetaldehyde. The acetaldehyde is then disproportionated to acetyl-CoA and ethanol and the acetyl-CoA is converted to acetate and ATP by genes within the microcompartment (Dde_3283, Dde_3279, Dde_3267, and Dde_3276 in Figure 2). The ethanol probably diffuses to the cytoplasm and is utilized as under ethanol/sulfate conditions, which explains why the cytoplasmic aldehyde:ferredoxin oxidoreductase (Dde_2460) is important for fitness on choline/sulfate (fitness = −0.79) as well as on ethanol.

*Lactate.* Based on our fitness data, the major lactate dehydrogenase in most of our lactate/sulfate or lactate/sulfite experiments seems to be Dde_3239:Dde_3240 (Figure 2). Dde_3244:Dde_3245 was also important for fitness in a few of the lactate experiments. A potential lactate dehydrogenase subunit (Dde_1842) and another lactate dehydrogenase (Dde_1085:Dde_1087) had little phenotype (Figure S1).

Although most of our experiments were conducted with mixed D,L-lactate, we performed one experiment each with 10 mM D-lactate or 10 mM L-lactate as the electron donor and 50 mM sulfate as the electron acceptor. The fitness profiles were very similar, with a linear (Pearson) correlation of 0.90. The most prominent differences in fitness were for a L-lactate permease (Dde_3238) and a nearby transcriptional regulator (Dde_3234), both of which were more important for fitness in L-lactate than in D-lactate (-0.26 vs. +0.41 and −0.68 vs. 0.0). In *D. vulgaris* Hildenborough, this regulator (DVU3023) binds upstream of and probably activates the expression of the permease (DVU3026) (Rajeev et al., 2011).

We observed secretion of succinate to 1.3–1.8 mM during growth in a defined medium with 60 mM lactate and 30 mM sulfate (Dataset S5). Keller et al. (2014) reported accumulation of succinate to 0.7 mM under similar conditions. *D. alaskensis* G20 can ferment pyruvate to acetate and succinate (Meyer et al., 2014) and apparently a similar metabolism takes place in the presence of lactate (which is oxidized to pyruvate) and sulfate.

*Fumarate and malate.* Fumarate and malate can be interconverted by fumarase (Dde_1254: Dde_1255). This enzyme is important for growth on either malate or fumarate with sulfate, but not under the other energetic conditions that we tested (Figure 2). The involvement of fumarase in malate utilization suggests that malate is being reduced to succinate as well as being oxidized to pyruvate. Specifically, malate would be oxidized by the decarboxylating malate dehydrogenase (Dde_1253), which would release pyruvate and reduced NADPH, while fumarate would be reduced to succinate by fumarate reductase, which would oxidize menaquinol. Unfortunately, we lack fitness data for the malate dehydrogenase, but fumarate reductase is important for fitness on both malate and fumarate but not in most other energetic conditions (Figure 2).

To verify that *D. alaskensis* G20 reduces fumarate and malate in the presence of sulfate, we measured the concentration of succinate in the media during growth in a defined medium with 10 mM fumarate or malate and 50 mM sulfate. During growth with fumarate and sulfate, we observed a “succinate burst” with a peak concentration of 2.3 mM during mid log phase (at 46 hours). During growth with malate and sulfate, we observed a much smaller release of succinate, to 0.2 mM. In both cases, the succinate disappeared after further growth. In the absence of sulfate, *D. alaskensis* G20 can ferment fumarate to acetate and succinate (Keller et al., 2014), and apparently this also occurs (at least temporarily) when sulfate is present.

The released succinate is probably reoxidized by fumarate reductase operating in reverse, i.e., succinate + menaquinone → fumarate + menaquinol. In other Desulfovibrio strains, succinate oxidation has been observed and seems to depend on the proton gradient (Zaunmiiller et al., 2006). The proton gradient should be required, as succinate oxidation with menaquinone is thermodynamically unfavorable. If succinate oxidation utilizes the proton gradient, then one might expect that the fumarate reductase reaction would create a proton gradient. However, the fumarate reductase of *D. alaskensis* G20 is related to the quinol:fumarate reductase of *Wolinella succinogenes,* which appears not to create a proton gradient when reducing fumarate. The *W. succinogenes* enzyme can nevertheless oxidize succinate *in vitro,* with roughly one proton moved per pair of electrons moved (Lancaster, 2013). So, we cannot determine whether fumarate reduction in *D. alaskensis* G20 leads to a proton gradient.

*Ethanol.* During ethanol oxidation, the ethanol is probably oxidized to acetaldehyde by one of two alcohol dehydrogenases (Dde_3523 or Dde_3534). This is expected to yield reduced NADH. Both of these genes have modest fitness defects that are specific to growth on ethanol (average fitness of −0.53 and −0.30), so they may be partially redundant. (This might also explain why they are not important for fitness on choline/sulfate.) The acetaldehyde would then be oxidized in the cytoplasm by acetaldehyde:ferredoxin oxidoreductase (Dde_2460, in Figure 2). This enzyme family yields acetate, not acetyl-CoA, as the product, so there is no opportunity for substrate-level phosphorylation.

*Interconversion of formate and hydrogen.* None of the four formate dehydrogenases were important for fitness during growth with formate as the electron donor, sulfate as the electron acceptor, and acetate as the carbon source (Figure S1). Instead, we found that the periplasmic [NiFeSe] hydrogenase (*hysAB*, Dde_2135:Dde_2134), was important for fitness, as were genes that are involved in its maturation (Figure 2). These results suggested that hydrogen, which is present in our anaerobic chamber and hence in the headspace of the hungate tubes, might be utilized instead of formate. However, when we tested control cultures with acetate but no formate added, no growth was observed. In contrast, the addition of both formate and acetate allowed significant growth: in defined media, the OD_600_ rose from 0.02 at inoculation to above 0.3. We then tested the growth of individual strains that had insertions in [NiFeSe] hydrogenase (*hysAB*); in *fdhAB* (Dde_0717:Dde_0718), which is the most highly expressed of the periplasmic formate dehydrogenases (Meyer et al., 2014); or in formate:hydrogen lyase (fhl). When grown individually, mutants of either *hysAB* or *fdhAB* showed little growth on formate but had normal growth on lactate/sulfate medium, while the *fhl* mutants grew about as well as the parent strain in either condition (Figure 3A & 3B).

The requirement for the [NiFeSe] hydrogenase suggests that formate is converted to hydrogen. Hydrogen release would also explain why the major formate dehydrogenase lacks a phenotype in the pooled assay, even though the mutant strain cannot grow in isolation.

It also appears that hydrogen is converted to formate. Formate was absent from our defined media (it was not intended to be present, and empirically its concentration was under 0.01 mM) but was present in the media during growth. In most growth conditions, formate was present at about 0.2 mM, but during growth with hydrogen as the electron donor, up to 0.6 mM formate was observed (Dataset S5).

If formate and hydrogen are interconverted, then the fitness patterns for the two electron donors should be very similar. Indeed, a number of energy-related genes were important for fitness on formate/sulfate but not on lactate/sulfate-these included the electron transport complexes Qrc and Rnf and genes for molybdopterin synthesis (Figure 3C). All of these genes were important during hydrogen utilization as well (Figure 3D). The role of Qrc and Rnf will be discussed in a later section. Molybdopterin synthesis is expected to be important for formate utilization because molybdopterin is part of the molybdenum or tungsten cofactor of the formate dehydrogenases. (Molybdopterin is also required for the activity of aldehyde:ferredoxin oxidoreductase or thiosulfate reductase, but these activities are probably not relevant under these conditions.) The mild loss of fitness for molybdopterin synthesis genes in the formate/sulfate fitness experiments (mean fitness = −0.47, *P <* 0.001, *t* test) suggests that cross-feeding of hydrogen does not fully make up for the inability to use formate in the pooled assay. Conversely, the molybdopterin synthesis genes may be slightly important for fitness during growth on hydrogen but not lactate (mean fitness = −0.27 versus +0.09, *P* < 0.003, paired *t* test). This is consistent with the conversion of hydrogen to formate.

We do not expect that the interconversion of formate and hydrogen would lead to a proton gradient. The major electron partner for both the periplasmic hydrogenases and formate dehydrogenases is probably Tp1-c_3_ (Pereira et al., 1998; Venceslau et al., 2010). (All of the periplasmic formate dehydrogenases in the *D. alaskensis* G20 genome are of the fdhAB type, without an associated cytochrome c_3_ subunit.) Another potential periplasmic electron carrier might be cytochrome c_553_ (Dde_1821), but mutants in this gene have little phenotype in any of our conditions (Figure S2), its only established role is as an electron donor for cytochrome c oxidase (Lamrabet et al., 2011), and it has a high redox potential (E^0ʹ^ = +0.02 V, Bianco et al. (1982)), which would prevent it from participating in sulfate reduction. If both hydrogen and formate are oxidized in the periplasm to reduce Tp1-c_3_, then it is hard to see how interconverting hydrogen and formate could yield energy.

**Figure S2.**
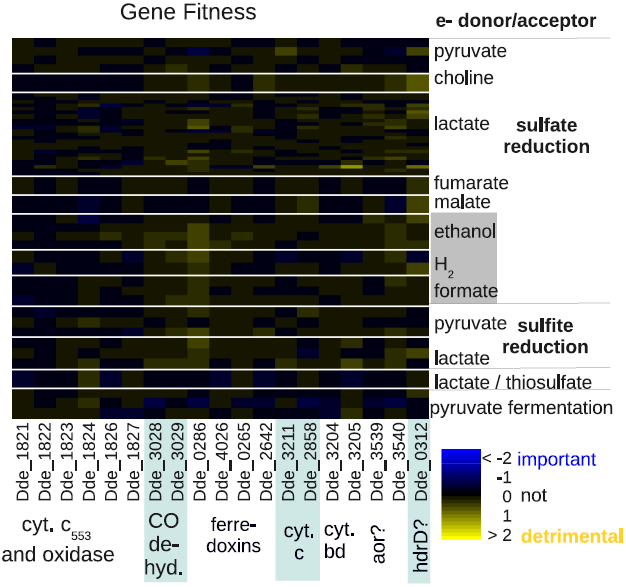
Heatmap of fitness data for electron transport genes that are not important for fitness in any of 12 energetic conditions.

Instead, we propose that utilizing formate requires converting some of it to hydrogen so that the hydrogen can diffuse to the cytoplasm and be reoxidized there. This would result in reduced ferredoxin that can be utilized by Rnf (discussed below). Reduced ferredoxin might also be necessary for the reduction of APS reductase (via a confurcation with Qmo). In our model, there is no other way for electrons from formate to reach ferredoxin (Figure 1D).

*Hydrogen oxidation.* During the oxidation of hydrogen, we did not observe a strong phenotype for any of the hydrogenases (Figure 2). The subunits of the [NiFeSe] hydrogenase had an average fitness of −0.39, and the subunits of the periplasmic [FeFe] hydrogenase had an average fitness of −0.06. We also grew individual mutants in the NiFeSe hydrogenase with hydrogen as the electron donor, and did not observe a growth defect (the maximum OD_600_ was 0.35–0.36 for the mutants and 0.36–0.38 for the parent strain). In contrast, a previous study found that a mutant in the periplasmic [FeFe] hydrogenase (*hydB*) had much reduced growth on hydrogen/sulfate media (Li et al., 2009). The difference might arise because our media contained added selenium, which would favor the expression of the [NiFeSe] hydrogenase over the [FeFe] hydrogenase (Valente et al., 2006), or because the level of hydrogen was lower in our study and the [FeFe] hydrogenase is a low-affinity hydrogenase (Caffrey et al., 2007). Overall, we propose that the periplasmic hydrogenases are redundant under our growth conditions. Another possibility is that hydrogen is primarily oxidized by the cytoplasmic [FeFe] hydrogenase (which we lack data for), although the observations of Li et al. (2009) suggest that periplasmic hydrogen oxidation is important.

## No evidence of energy conservation by molecular cycling

As discussed above, we found that the periplasmic [NiFeSe] hydrogenase was important during the oxidation of formate, but we do not expect that this contributes to energy conservation. In fitness experiments with sulfate and no added hydrogen or formate, hydrogenases or formate dehydrogenases were not important for fitness (Figures 2 & S1). And as mentioned above, individual growth assays confirmed that mutants in the major periplasmic formate dehydrogenase or hydrogenase grew as well as the parent strain under lactate/sulfate conditions (Figure 3B). In contrast, in *D. vulgaris* Hildenborough, formate dehydrogenases are important for growth on lactate/sulfate media (da Silva et al., 2013). It has also been proposed that carbon monoxide cycling plays a role in energy conservation in *D. vulgaris* Hildenborough (Voordouw, 2002). *D. alaskensis* G20 encodes a CO dehydrogenase (Dde_3028:Dde_3029) but it is not important for fitness under any of our growth conditions (Figure S2).

Overall, our data suggests that in *D. alaskensis* G20, molecular cycling of hydrogen, formate, or carbon monoxide is not important for energy conservation during sulfate reduction. Although we do not have fitness data for two key parts of the hydrogen cycling model, the cytoplasmic [FeFe] hydrogenase or the periplasmic electron carrier Tp1-c_3_ (*cycA*), previous studies found that *cycA* mutants of *D. alaskensis* G20 grew about as well as the parent strain in lactate/sulfate media (Rapp-Giles et al., 2000; Keller et al., 2014), albeit with increased secretion of formate (Li et al., 2009), which is an alternate destination for electrons from Tp1-c_3_.

## Utilization of electron acceptors

*Sulfate.* While we were measuring metabolites to understand growth with sulfate as the electron acceptor and different electron donors, we observed that *D. alaskensis* G20 does not always reduce all of the sulfate to sulfide. During growth with H_2_ as the electron donor, we detected the release of 5–7 mM thiosulfate, which corresponds to 10 mM or more of sulfur atoms, or about a third of the sulfur that was originally added (as 30 mM sulfate). During growth with 60 mM lactate as the electron donor and 30 mM sulfate as the electron acceptor, about 1 mM thiosulfate was formed. These results were not expected, as sulfate-reducing bacteria are believed to reduce sulfate all the way to sulfide (Rabus et al., 2013) or to release thiosulfate transiently (Vaïnshteïn et al., 1979; Fitz and Cypionka, 1989). The highest reported level of thiosulfate release by sulfate-reducing bacteria that we are aware of is just 0.4 mM (Sass et al., 1992). However, dissimilatory sulfite reductase does release thiosulfate *in vitro* when electrons are limiting: it appears that sulfite can attack S^0^ while it is bound by DsrAB to release thiosulfate (Parey et al., 2010). Although DsrAB could later reduce the reduced thiosulfate to sulfide, the reduction of thiosulfate by hydrogen is thermodynamically not favorable (Δ*G*^0′^ = −4 kJ/mol, but this corresponds to an unrealistically high 1 atm of H_2_, Cypionka (1995)).

*Sulfite.* We did not find strong phenotypes during growth with sulfite as the electron acceptor (Figures 2 & S1), probably because the genes involved in sulfite reduction are also essential during growth on sulfate.

*Thiosulfate.* Unexpectedly, the putative thiosulfate reductase PhsAB (Dde_1076:Dde_1075) did not have strong phenotypes during growth on lactate/thiosulfate (fitness of −0.79 and −0.30, respectively). The molybdopterin synthesis genes, which are required to produce the molybdenum cofactor of thiosulfate reductase, also had moderate phenotypes (mean fitness −0.75, *P* = 0.005, *t* test). The high activity of DsrAB for thiosulfate reduction (Parey et al., 2010) might allow these mutants to grow.

*Pyruvate fermentation. D. alaskensis* G20 can disproportionate pyruvate to acetate and succinate, via reduction of pyruvate to malate and then succinate (Meyer et al., 2014). Fumarate reductase was important for fitness during pyruvate fermentation, but to a varying extent, with a mean fitness −0.92 in one experiment and −0.51 and −0.35 in the others. During the experiment in which fumarate reductase was most important, fumarase was also important (fitness = −1.0). The importance of fumarase and fumarate reductase is consistent with a previous measurement of the fitness of these pools of mutants during pyruvate fermentation (Meyer et al., 2014). However, the electron transport complexes Hdr/flox-1 and Rnf were important for fitness in the previous study but not in ours (mean fitness = +0.03 and +0.58, respectively). Whereas our experiments were conducted in sealed hungate tubes, the previous study used a bioreactor with slow flushing of the headspace, which would remove H_2_ from the system; this might account for the difference.

## Roles of electron transport complexes in energy conservation

Among the electron transport complexes, we observed clear phenotypes for Qrc, Rnf, Hdr/flox-1, NfnAB-2, Nox, Tmc, and Hmc (Figure 4).

*Qrc.* Our fitness data for Qrc (Dde_2932:Dde_2935) is consistent with previous reports that it is important for hydrogen or formate oxidation but not for growth on lactate/sulfate media (Li et al., 2009; Keller et al., 2014). Qrc is a Tp1-c_3_:menaquinone oxidoreductase and in combination with Qmo, it is proposed to pump protons while feeding electrons from periplasmic Tp1-c_3_ into sulfate reduction. This explains why it is important for the utilization of hydrogen or formate, which are oxidized in the periplasm. But we also observed that Qrc was important for the utilization of fumarate, malate, and ethanol, which are oxidized in the cytoplasm, and Qrc had moderate phenotypes in a few of the lactate/sulfate experiments (lowest mean fitness of −1.15).

Qrc is probably important with these electron donors because it is part of the path to menaquinone. In particular, it appears that Tp1-c_3_ is required for sulfate reduction with pyruvate as the electron donor because electrons flow via Qrc from Tp1-c_3_ to reduce menaquinone, which is required by Qmo (Keller et al., 2014). Tp1-c_3_ is not required in the presence of lactate, presumably because lactate dehydrogenase provides reduced menaquinone. The requirement for oxidizing Tp1-c_3_ or reducing menaquinone explains all of the fitness data for Qrc, except that we found only a mild defect for *qrc* mutants with pyruvate as the electron donor and sulfate as the electron acceptor (mean fitness of −0.90 to −0.16). In contrast, when a *qrcA* mutant is grown individually in pyruvate/sulfate media, it grows poorly, with a greatly extended lag (K. L. Keller and J. D. Wall, personal communication). To explain this discrepancy, we note that *qrc* mutants grow well, relative to the parent strain, by pyruvate fermentation (Figure 4, and Meyer et al. (2014)). So, we propose that in the pooled assay, the *qrc* mutants are able to grow by fermenting pyruvate. Although growth by pyruvate fermentation is normally much slower than growth by pyruvate oxidation (Keller et al., 2014), the *qrc* mutants would benefit from the removal of hydrogen (or other end products) by other strains that are reducing sulfate.

*Rnf.* The ion-pumping ferredoxin:NADH oxidoreductase Rnf (Dde_0581:Dde_0587, rn-fCDGEABF) was important for growth with sulfate as the electron acceptor and malate, fumarate, ethanol, hydrogen, or formate as the electron donor (Figure 4). Growth assays with individual mutants confirmed that Rnf mutants grew little or not at all in most of these conditions (Figure 5). (The exceptions were that a mutant in *rnfF* sometimes reached a high yield after an extended lag, and we did not test growth in malate/sulfate media.) These observations are consistent with a previous report that Rnf is required for the utilization of hydrogen or formate by *D. alaskensis* G20 (L. R. Krumholz *et al.* 2011, DOE Hydrogen and Cells Program Annual Progress Report). In contrast, the decaheme cytochrome (*dhcA*), which is cotranscribed with rnfCDGEABF and is the first gene in the operon, is not important for fitness except in one pyruvate fermentation experiment (Figure 4). The function of DhcA is not known, but our data suggests that it is not involved in electron transport by Rnf, at least not in our growth conditions.

**Figure 5.**
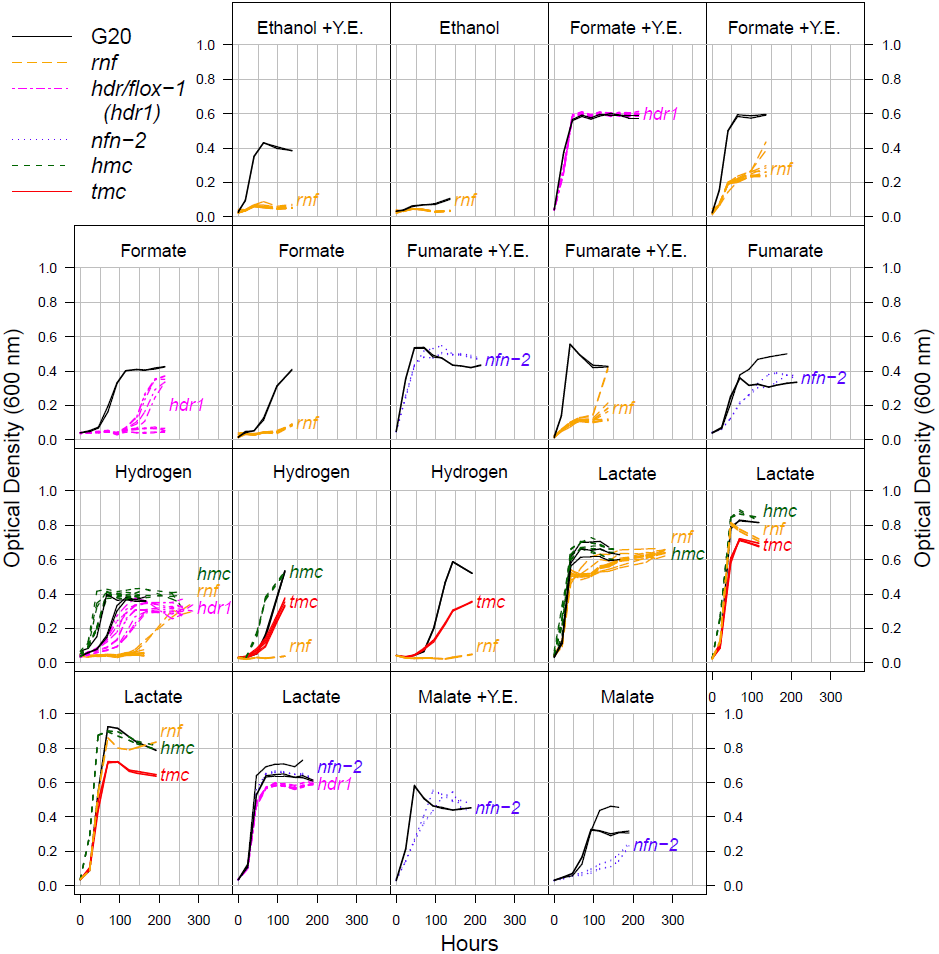
Growth of mutants in electron transport complexes Rnf, Hdr/flox-1, Nfn-2, Hmc, and Tmc, with sulfate as the electron acceptor and with a variety of electron donors. Growth of the parent strain (G20) is shown for comparison. Y.E. is short for yeast extract. Experiments done on different days are graphed separately. There are transposon insertions within more than one member of each complex, and there are 2-4 replicates for each mutant and condition, except for floxD-1 (Dde 1210) growing on hydrogen.

As Rnf can create an ion gradient, the most obvious explanation for its phenotypes is that it is involved in energy conservation. Indeed, it is required for growth with all of the electron donors that do not allow for substrate-level phosphorylation and for which the requirement for ion pumping might be greatest (ethanol, hydrogen, and formate). Also, all of the electron donors that it is important for are expected to lead to reduced ferredoxin. (Malate and fumarate are oxidized to pyruvate, which is a substrate for pyruvate:ferredoxin oxidoreductase. Ethanol is oxidized to acetaldehyde, which is a substrate for acetaldehyde:ferredoxin oxidoreductase. Hydrogen is a substrate for a cytoplasmic [FeFe] ferredoxin hydrogenase. Finally, formate seems to yield cytoplasmic hydrogen as discussed above.) Another circumstantial piece of evidence for Rnf’s role in energy conservation is that in *D. alaskensis* G20 and other Desulfovibrio species, Rnf and many genes that are involved in sulfate reduction are coregulated by the redox-responsive regulator Rex (Ravcheev et al., 2012).

A second possibility is that Rnf operates in reverse, to produce reduced ferredoxin that is otherwise unavailable. This seems unlikely because of the energetic cost and because of the expected availability of reduced ferredoxin on these electron donors. A related question is which ferredoxin is oxidized (or reduced) by Rnf. We expect that it reduces ferredoxin I (Dde_3775), which is the major cytoplasmic electron carrier (Ogata et al., 1988) and may be essential in Desulfovibrios (Fels et al., 2013; Kuehl et al., 2014). Other ferredoxins are not important for fitness in these conditions (Figure S2).

Third, Rnf could be involved in cofactor synthesis. For example, in *D. vulgaris* Hildenborough, Rnf is required for nitrogen fixation (Keller and Wall, 2011), presumably because it reduces a nitrogenase-specific ferredoxin, as in *Rhodobacter capsulatus,* where Rnf was first described (Schmehl et al., 1993). However, the genome of *D. alaskensis* G20 does not include genes for nitrogen fixation, and no other role for Rnf in cofactor synthesis has been reported (Biegel et al., 2011).

Finally, some homologs of Rnf are involved in signalling. For example, in *E. coli*, Rnf is known as Rsx: it reduces the FeS cluster of the SoxR transcriptional activator to eliminate its activity in the absence of oxidizing stresses (Koo et al., 2003). *D. alaskensis* G20 contains a potential soxR-like regulator (Dde_2633) but our fitness data does not suggest a relationship between SoxR and Rnf (the correlation of fitness patterns is 0.05, *P* > 0.5, *n* = 49) and SoxR did not have strong phenotypes in our energetic conditions (the range of fitness values was −0.5 to +0.7). Also, it is not obvious why increasing the response to a redox stress would eliminate growth under a subset of energetic conditions.

It is also interesting that the genomes of many sulfate-reducing bacteria encode Rnf but some Desulfovibrio species do not (Pereira et al., 2011). The Desulfovibrio genomes that do not encode Rnf do encode a proton-pumping hydrogenase (Ech and/or Coo), which can create an ion gradient while moving electrons from ferredoxin to hydrogen. The *D. alaskensis* G20 genome does not encode Ech or Coo. Thus, different members of the Desulfovibrio genus may use different systems to create an ion gradient while transferring electrons from reduced ferredoxin.

*Hdr/flox-1.* A recent review proposed that a cluster of heterodisulfide-reductase-like (hdr) genes with flavin oxidoreductase (flox) genes should be named Hdr/flox (Pereira et al., 2011) . Although this complex has not been studied experimentally, it was proposed to perform an electron bifurcation from NADH to a ferredoxin and a heterodisulfide electron carrier such as DsrC. The genome of *D. alaskensis* G20 encodes two paralogous Hdr/flox operons, which we will term Hdr/flox-1 (Dde_1207:Dde_1213) and Hdr/flox-2 (Dde_3524: Dde_3530). Despite the potential redundancy of these operons, Hdr/flox-1 was important for growth on formate (mean fitness −1.2, *P* < 10^−5^, *t* test). Hdr/flox-1 also had mild fitness defects on hydrogen (mean fitness −0.54, *P* < 10^−5^, *t* test) and ethanol (mean fitness −0.44, *P* < 0.0001, *t* test). Growth curves for individual mutants in *hdr/flox-1* confirmed that they had a severe growth defect in defined formate/acetate/sulfate media and a modest growth defect in defined acetate/sulfate media with added hydrogen, but they had little or no reduction in growth in defined lactate/sulfate media or in a rich formate medium (Figure 5).

As Rnf is required for growth on formate, hydrogen, and ethnol, we propose that Hdr/flox-1 converts NADH from Rnf back to ferredoxin to allow additional ion pumping, while feeding electrons into the sulfite reduction pathway. If Rnf runs twice for each iteration of Hdr/flox, and Rnf pumps one ion per pair of electrons transferred, then the overall reaction would be Fd^2−^ + DsrC_*ox*_ + 2 ion_*cytoplasm*_ → Fd^0^ + DsrC_*red*_ + 2 ion_*periplasm*_.

*NfnAB-2.* The genome of *D. alaskensis* G20 enodes two paralogous operons for the electron-bifurcating transhydrogenase NfnAB, which we will term NfnAB-1 (Dde_3635:Dde_3636) and NfnAB-2 (Dde_1250:Dde_1251). NfnAB-2 had a mild fitness defect on malate/sulfate and fumarate/sulfate experiments (average fitness of −0.76 and −0.55). Growth curves with individual mutants confirmed that *nfn-2* mutants grew more slowly than the parent strain in malate/sulfate and fumarate/sulfate media, whether or not yeast extract was added (Figure 5). As the oxidation of malate (or fumarate) yields NADPH, we propose that Nfn is oxidizing NADPH and reducing ferredoxin and NAD^+^. This is an energy conserving mechanism because the reduced ferredoxin could yield an ion gradient via Rnf.

In other conditions, Nfn might run in the opposite direction, to use the energy in low-potential ferredoxin to drive electrons to NADPH and maintain a high NADPH/NADP^+^ ratio. In fact, we do not know of another mechanism by which Desulfovibrios could maintain a high NADPH/NADP^+^ ratio. However, the phenotypes for NfnAB-2 were observed in conditions where the oxidation of malate should generate reduced NADPH, and in the presence of yeast extract, which would minimize the need for NADPH for biosynthetic reactions.

*Nox.* NADH oxidase (Nox, Dde_0374) can reduce oxygen to hydrogen peroxide (Chen et al., 1994), and there are varying reports as to whether it interacts with and transfers electrons to APS reductase (Chen et al., 1994; Chhabra et al., 2011). Thus, it is not clear whether Nox is involved in the transfer of electrons from NADH into the sulfate reduction pathway. We found that Nox was important for fitness in many of our energetic conditions, regardless of whether sulfate was present (Figure 4). Thus, although Nox seems to have an important role, it does not seem to be specific to sulfate reduction.

The phenotypes of Nox were often not consistent across similar experiments, which could indicate that the data for this gene is not reliable. However, Nox was strongly co-fit with two uncharacterized cotranscribed genes, Dde_3773:Dde_3772, across our data set (r = 0.80 and 0.79, respectively; these were the two most co-fit genes). Such a correlation is very unlikely to occur by chance (uncorrected *P <* 10^−11^, or *P* = 10^−8^ after correcting for multiple testing across all of the genes in our data set). Finding two genes in the same operon as the most cofit genes also suggests that the fitness pattern is genuine. We also observed that Nox is very important for surviving oxygen stress (mean fitness = −3.8), which is consistent with its biochemical function *in vitro.* We speculate that Nox is important for resisting redox stresses that are present at variable levels in our experiments.

*Hmc and Tmc.* Hmc and Tmc are multi-subunit transmembrane electron transfer complexes that are believed to exchange electrons with Tp1-c_3_ and transfer them across the membrane (Pereira et al., 1998, 2006; Quintas et al., 2013). In both cases, the redox partner in the cytoplasm is not known, but DsrC or ferredoxin have been suggested (Venceslau et al., 2014; Walker et al., 2009). Some of the subunits of the two complexes are homologous to each other, and Tmc might be a simplified form of Hmc (Pereira et al., 2011). In particular, the Hmc complex, but not the Tmc complex, contains an NrfD-like subunit (HmcC). Other members of the NrfD family are proposed to be menaquinone-interacting proton pumps (Jormakka et al., 2008).

We found that Tmc was important for hydrogen oxidation (mean fitness = −1.6) but Hmc was not (mean fitness = +0.1). We observed other mild phenotypes for both complexes, which were consistent across mutant strains in each complex but were not consistent across similar conditions, so they are difficult to interpret (Text S2). By growing individual strains, we confirmed that *hmc* mutants have a growth advantage on hydrogen/sulfate, with the lag reduced by almost one day relative to the parent strain, while *tmc* mutant strains had slower growth (Figure 5).

Our genetic data for *D. alaskensis* G20 is consistent with the observation that Tmc is reduced by Tp1-c_3_ and hydrogenases *in vitro* (Pereira et al., 2006). During hydrogen oxidation, Qrc is also important for fitness, and Qrc sends electrons from Tp1-c_3_ to menaquinone, from which they probably go to Qmo and ultimately reduce APS. Another path from Tp1-c_3_ to APS would seem redundant, so we propose that Tmc is necessary because it sends electrons from Tp1-c_3_ towards the sulfite reduction pathway (i.e., DsrC). We also note that DsrMKJOP is believed not to accept electrons from Tp1-c_3_ (Pires et al., 2006), which explains why another path from Tp1-c_3_ to DsrC is needed.

We measured the release of hydrogen during growth of the parent strain and mutants in Hmc and Tmc in defined lactate/sulfate media. We observed an increased maximum level of hydrogen in *tmc* mutants (1,120–2,192 ppm versus 641–819 ppm for G20) and the burst persisted for much longer: for example, at 140 hours, *tmc* mutants had 606–879 ppm of H_2_ remaining while the other strains had a maximum of 194 ppm remaining (Figure S3A). This shows that Tmc is involved in the utilization of hydrogen during growth on lactate as well.

**Figure S3.**
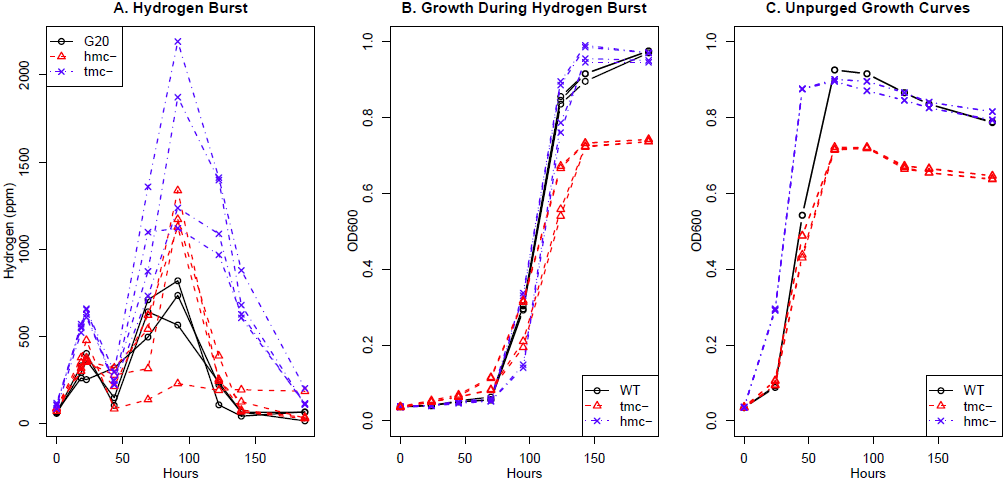
A. Hydrogen levels in G20, *hmc* mutant, and *tmc* mutant strains growing in defined media with 60 mM lactate and 30 mM sulfate. The head space was purged with 80:20 N_2_:CO_2_ at the start of the experiment. The *hmc* mutants have insertions in *hmcE* or *hmcF;* the *tmc* mutants have insertions in *tmcB* or *tmcC;* and there are two replicates for each strain. B. Growth during the hydrogen experiment. C. Growth curves performed on the same day but without purging the headspace. Again, there are two replicates for each strain.

Finally, we observed that Tmc is important for resisting tetrakis-hydroxymethyl phosphonium sulfate (THPS) stress during growth in lactate/sulfate media, with a mean fitness −1.8. (These experiments were conducted in “MO” media with added vitamins and the Tmc mutants were not sick in THPS-free lactate/sulfate fitness experiments that were performed on the same days.) THPS is a biocide that is effective against sulfate-reducing bacteria and gene expression data suggests that it targets their energy metabolism (Lee et al., 2010). The phenotype for the Tmc complex confirms that THPS affects the energy metabolism of *D. alaskensis* G20.

As far as we know, mutants in Tmc have not been studied before, but in both *D. vulgaris* Hildenborough and *D. alaskensis* G20, Hmc is important for syntrophy with a methanogen, during which the Desulfovibro ferments lactate or pyruvate to acetate and either CO_2_ and H_2_ or formate, while the methanogen consumes the H_2_ or formate so that the fermentation becomes energetically favorable (Walker et al., 2009; Li et al., 2011; Meyer et al., 2013). This suggests that Hmc sends electrons from a lower-potential donor in the cytoplasm such as ferredoxin (which is produced by oxidizing pyruvate) to the periplasmic Tp1-c_3_. If Hmc sends electrons from ferredoxin to Tp1-c_3_, then there would be sufficient energy to drive the export of 1 or 2 protons per electron pair. In *D. vulgaris* Hildenborough, Hmc is also important for growth on plates with lactate as the electron donor and no reductant in the media (Dolla et al., 2000), which was explained by proposing that Hmc is required to reduce the redox potential of the media (Dolla et al., 2000), which again suggests that Hmc is sending electrons from the cytoplasm to the periplasm. Consistent with this model, we found that in *D. alaskensis* G20, Hmc was important for fitness during growth on lactate/sulfate agar plates (mean fitness = −2.2) and for surviving oxygen stress (mean fitness = −2.9). We also note that an *hmc* mutant strain of *D. vulgaris* Hildenborough showed a roughly 30% reduction in the rate of hydrogen utilization (Dolla et al., 2000). This contrasts to our finding for *D. alaskensis* G20 but could relate to hydrogen utilization in the cytoplasm by *D. vulgaris* Hildenborough, which would be coupled to ferredoxin reduction.

Another recent study reported that the “Hmc” complex of *D. piger* GOR1 was important for hydrogen utilization (Rey et al., 2013), but this complex was misannotated. The genome does not contain *hmcE* or *hmcF*, and the homolog of *hmcA* is shortened; overall, the “Hmc” operon is very similar to the Nhc operon of *D. desulfuricans* ATCC 27774, whose first gene encodes a nine-heme cytochrome rather than the 16-heme cytochrome *hmcA* (Matias et al., 1999; Saraiva et al., 2001). Hence, in both *D. piger* GOR1 and *D. desulfuricans* G20, a simplified form of Hmc is required for hydrogen utilization.

In summary, we showed that Tmc is important for utilizing hydrogen as an electron donor and for consuming hydrogen that was previously released during growth on lactate. Thus, it appears that Tmc transfers electrons from Tp1-c_3_ to the sulfite reduction pathway, perhaps to DsrC. In contrast, Hmc’s activity is detrimental during growth with hydrogen as the electron donor, and we propose that it transfers electrons from ferredoxin to Tp1-c_3_ and creates an ion gradient.

## No genetic evidence for other routes of electron transfer

Besides the genes discussed so far, *D. alaskensis* contains numerous genes that have been proposed to play a role in electron transport and/or sulfate reduction (Pereira et al., 2011). Mutants in most of these genes showed little phenotype in our fitness experiments (Figure S2). The genes without phenotypes included a variety of putative electron carriers, such as the alternate cytoplasmic ferredoxin (ferredoxin II, Dde_0286) and other putative ferredoxins; periplasmic split-Soret cytochrome c (Dde_3211 or Dde_0653); and a periplasmic c_554_-like cytochrome (Dde_2858). We cannot be sure that these proteins are not carrying a significant flow of electrons, as a strain that lacks one route of electron transfer might be able to compensate by using another redundant pathway. But the simplest interpretation of our results is that these redox proteins are not important under any of our growth conditions.

## Conclusions

Despite the large number of electron carriers and electron transfer complexes in the genome of *D. alaskensis* G20, we identified phenotypes for many electron transfer genes under a subset of energetic conditions. These confirmed the expected path of electrons from choline, lactate, fumarate, malate, ethanol, or formate into central energy metabolism. We also showed that Tmc is involved in the oxidation of hydrogen by *D. alaskensis* G20, probably by moving electrons from periplasmic Type 1 cytochrome c_3_ to the cytoplasmic sulfate reduction pathway (perhaps via DsrC). In contrast, the Hmc complex was detrimental to growth with hydrogen as the electron donor and was important for survival in the presence of oxygen.

We found little evidence for energy conservation via the cycling of hydrogen, formate, or carbon monoxide. Instead, the phenotypes for mutants in Rnf, Hdr/flox-1, and NfnAB-2 suggest that these complexes are involved in energy conservation by pumping ions or by electron bifurcations that allow the reduction of an electron carrier with a low redox potential, such as ferredoxin. However, Hdr/flox has never been studied biochemically in any organism, and it will be important to identify its redox partners and to determine whether it actually performs an electron bifurcation. Surprisingly, we found that the periplasmic [NiFeSe] hydrogenase (HysAB) is important for formate utilization. The resulting hydrogen could diffuse to the cytoplasm, where the cytoplasmic hydrogenase would reduce ferredoxin and allow ion pumping by Rnf.

Our genetic approach is ill suited to studying genes that are essential for sulfate reduction. Another limitation of our approach is genetic redundancy. Although Hdr/flox and Nfn have paralogs in *D. alaskensis* G20, they are present as a single copy in *D. vulgaris* Hildenborough and *D. vulgaris* Miyazaki F, and we hope to get a clearer picture of their roles by studying mutants of those organisms. Another potential way to overcome genetic redundancy would be to study double mutants.

Finally, a long-standing mystery in the energetics of Desulfovibrios has been the role of NAD(P)H. Reduced NADH is probably formed by ethanol dehydrogenase, and reduced NADPH is probably formed by malate dehydrogenase, but the path for electrons from NAD(P)H to sulfate reduction has not been clear. Our data suggest that electron bifurcations by Hdr/flox and Nfn can allow electrons from NAD(P)H to feed into sulfate reduction.

## Methods

### Strains and growth conditions

*D. alaskensis* G20 was provided by Terry Hazen (University of Tennesse, Knoxville). The transposon mutants that were used for growth curve experiments or metabolite measurements were verified by streaking out single colonies and using colony PCR to verify that the transposon insertion was at the expected location. The mutants and primers are listed in Dataset S4.

Fitness experiments and growth curve experiments were conducted anaerobically in hungate tubes at 30°C. Media was prepared within a Coy anaerobic chamber with an atmosphere of about 2% H_2_, 5% CO_2_, and 93% N_2_. Although some H_2_ is present in all experiments, control experiments showed that it does not suffice to support growth.

Two base media formulations were used – “Hazen” and “MO” media. Hazen media was used for 16 fitness experiments; MO media was used for 33 fitness experiments and for all growth curves. Hazen base media contained 30 mM PIPES buffer at pH 7.2, 20 mM NH_4_Cl, other salts (see below), 0.625 mM nitriloacetic acid as a chelator, and 0.016 μM resazurin as a redox indicator. MO base media contained 30 mM Tris-HCl buffer at pH 7.2, 5 mM NH_4_Cl, other salts, and 0.12 mM EDTA as a chelator, but no redox indicator. For both base media, a reductant was usually included: for Hazen media, the reductant was usually 0.38 mM titanium citrate (15/16 experiments), while for MO media, the reductant was usually 1 mM Na_2_S (31/33 experiments).

For the two base media, the composition of the salts was similar, but trace metals were at roughly two-fold higher concentration in Hazen media than in MO media. For Hazen media, salts were added to a final concentration of 8 mM MgCl_2_, 0.6 mM CaCl_2_, 2.2 mM K_2_HPO_4_, and 62.5 *μ*m FeCl_2_, as well as trace metals: 31.25 *μ*m MnCl_2_, 16.25 *μ*m CoCl_2_, 18.75 *μ*m ZnCl_2_, 2.625 *μ*m Na_2_MoO_4_, 4 *μ*m H_3_BO_3_, 4.75 *μ*m NiSO_4_, 0.125 *μ*m CuCl_2_, 0.375 *μ*m Na_2_SeO_3_, and 0.25 *μ*m Na_2_WO_4_, For MO media, salts were added to a final concentration of 8 mM MgCl_2_, 0.6 mM CaCl_2_, 2 mM K_2_HPO_4_, and 60 *μ*m FeCl_2_, as well as trace metals: 15 *μ*m MnCl_2_, 7.8 *μ*m CoCl_2_, 9 *μ*m ZnCl_2_, 1.26 *μ*m Na_2_MaO_4_, 1.92 *μ*m H_3_BO_3_, 2.28 NiSO_4_, 0.06 *μ*m CuCl_2_, 0.21 *μ*m Na_2_SeO_3_, and 0.144 *μ*m Na_2_WO_4_.

To these base media, we added various electron donors and acceptors, 0.1% yeast extract (24/49 fitness experiments), and/or 1 ml/L of Thauer’s vitamin solution (22/49 fitness experiments; Brandis and Thauer (1981)). However, vitamins were not added for growth curve experiments.

If sulfate was the electron acceptor, it was added to a final concentration of either 15 mM (Hazen media), 30 mM (MO media), or 50 mM (when fumarate or malate were the electron donors). Other electron acceptors (sulfite or thiosulfate) were added at 10 mM. If lactate was the electron donor, it was usually at 60 mM (9/12 Hazen fitness experiments and 12/15 MO fitness experiments) but several fitness experiments used 10 mM or 15 mM. Choline was at 30 mM. The concentration of pyruvate was 20 mM (Hazen) or 60 mM (MO) if sulfate was the electron acceptor, 30 mM if sulfite was the electron acceptor, or 60 mM for pyruvate fermentation experiments. Ethanol was at 10 mM (Hazen) or 60 mM (MO). Fumarate or malate were at 10 mM. Formate was at 50 mM. Hydrogen gas was added by blowing a mix of hydrogen (80%) and CO_2_ (20%) through the culture for 2 minutes, either once or periodically (five times total). For formate and hydrogen experiments, acetate was also added as a carbon source, to a final concentration of 10 mM.

### Fitness experiments

Of the 49 energy-related fitness experiments analyzed in this paper, 43 are newly described here. Four experiments in lactate/sulfate media are described by Price et al. (2013). Two experiments with lactate/sulfate or choline/sulfate media are described by Kuehl et al. (2014). For complete metadata of all 49 energy-related fitness experiments, see Dataset S1. We also conducted 33 other fitness experiments: growth on agar plates with rich or defined lactate/sulfate medium; survival of oxygen stress in rich lactate/sulfate medium followed by outgrowth in a rich lactate/sulfate medium; and growth in lactate/sulfate media in the presence of various compounds or with heat stress at 42°C. These are also included in Dataset S1.

Fitness experiments were conducted and analyzed as described previously (Price et al. 2013). Briefly, for each of the two pools, we collected a sample at the start of the experiment (at inoculation) and at the end of growth. For each sample, we extracted genomic DNA and we used PCR to amplify the DNA barcodes from the “tag modules” that distinguish the strains within each pool (Oh et al., 2010). Each tag module contains an “uptag” and “downtag” barcode. For most of the fitness experiments, we amplified both the uptags and the downtags from each sample, mixed then together, hybridized them to an Affymetrix 16K TAG4 microarray, and scanned the microarray (Pierce et al., 2007). In these cases there were two microarrays per fitness experiment. For four of the fitness experiments, we amplified the uptags from one pool, the downtags from the other pool, and mixed these together, so that there was just one microarray for the fitness experiment.

To estimate the abundance of a strain, we averaged the log_2_ intensity across replicate spots and across probes for the uptag and downtag (if we amplified both tag modules from that sample). The fitness value for a strain is then the change in log_2_ intensity.

The fitness value for a gene is the average of fitness values for the relevant strain(s). This per-gene fitness value was normalized so that the mode of the distribution was zero. Then, we used smooth local regression (loess) to remove any effect of distance from the origin of replication on the fitness values. Such effects might be an artefact of variation in copy number across the chromosome in growing cells.

The per-strain and per-gene fitness values are included in Dataset S2 and Dataset S3, respectively. All of the fitness data is also available on MicrobesOnline.

### Concentrations of metabolites

To measure the concentrations of various compounds during growth, we grew the parent G20 strain in a defined MO media with no vitamins added and with malate (10 mM), fumarate (10 mM), acetate (10 mM), or lactate (60 mM) as the carbon source. Sulfate was the electron acceptor, at 50 mM for malate or fumarate experiments or 30 mM otherwise. If acetate was the carbon source, hydrogen gas was added as an electron donor at the beginning of the experiment as a 20% mix with CO_2_. Culture samples (0.5 ml) were taken roughly once per day until the culture stopped growing. For each metabolite sample, the sample was spun down at 14,000 × g for 10 minutes at room temperature and the supernatant was analyzed by ion chromatography and HPLC.

For ion chromatography (ICS-5000+, Thermo-Fisher Scientific), a 25 *μ*L aliquot of the supernatant was injected onto an IonPac AS11-HC Analytical Column (4 × 250 mm, ThermoFisher Scientific) equipped with a guard column (4 × 50 mm) of the same material. Pyruvate, formate, fumarate, lactate, sulfate, thiosulfate, phosphate, and chloride were eluted at 30°C by a gradient program of 0.2 mM sodium hydroxide for 6 min, then in 5 min to 5 mM, then in 16 min to 40 mM, at a flow rate of 2 mL/min, and detected by suppressed conductivity.

Another aliquot (20 *μ*L) of the supernatant was injected onto an Aminex HPX-87H column (7.8 300 mm, Bio-Rad) and malate and succinate were eluted at 50°C using 5 mM sulfuric acid at a flow rate of 0.6 mL/min and detected by refractive index. The identity of succinate was additionally confirmed by accurate mass spectrometry (6520 QTOF, Agilent Technologies).

The concentrations of metabolites, and the optical densities for the corresponding time points, are provided in Dataset S5.

### Hydrogen measurements

To measure the concentration of hydrogen during growth in lactate/sulfate media, we performed a similar growth experiment but with *hmc* and *tmc* mutants as well as with the parent G20 strain, and we purged the headspace with an 80:20 N_2_:CO_2_ mix at the beginning of the experiment. These experiments had a much longer lag phase than the other metabolite experiments (roughly 75 hours instead of 20 hours for the G20 strain) – apparently the absence of hydrogen in the headspace at the start of the experiments delayed growth (Figure S3B & S3C). Roughly once per day, 0.5 ml of the headspace was removed and analyzed.

The concentration of H_2_ gas was quantified using gas chromatography with a Bruker 450-refinery gas analyzer (Bruker Daltonics) equipped with a HayeSep N and a molecular sieve packed column coupled in series and kept at 50^°^C. and injected into a flow of 30 mL/min nitrogen, and hydrogen was detected by thermal-conductivity detection. A one-point calibration was performed by injecting 0.5 mL of a custom blend gas mix (prepared in nitrogen, Scott-Marrin Inc.).

## Supplementary Material

Dataset S1: Metadata for the fitness experiments. Energy-related experiments are given in the same order as in the heatmaps, followed by additional experiments. Tab-delimited file: http://morgannprice.org/G20energy/G20_energyfitness_experiments.xls

Dataset S2: Per-strain fitness values. Tab-delimited file: http://morgannprice.org/G20energy/G20_energy_strainfitness.xls

Dataset S3: Per-gene fitness values. Tab-delimited file: http://morgannprice.org/G20energy/G20_energy_gene_fitness.xls

Dataset S4: Mutants for follow-up studies and primers for confirming the transposon insertion location in each strain. Tab-delimited file: http://morgannprice.org/G20energy/Single_mutants.tab

Dataset S5: Concentrations (mM) of metabolites in the supernatant during growth of *D. alaskensis* G20 in defined media. Tab-delimited file: http://morgannprice.org/G20energy/G20_supernatant.concentrations.xls

## Text S1: Energy-related genes or complexes that are absent from the fitness data.

We lack data for genes that are essential for the activation of sulfate (sulfate adenyltransferase, adenylate kinase, and pyrophosphatase Dde_1778), the reduction of APS (APS reductase or qmoABC), or the reduction of sulfite (dsrABC or dsrMKJOP, although we have an insertion at the very 3’ end of dsrP). We also lack data for other genes or operons that are probably required for energy production on lactate/sulfate media, notably a cluster of genes for the oxidation of pyruvate to acetyl-CoA and conversion to acetate and ATP (Dde_3237, Dde_3241, and Dde_3242); ferredoxin I (Dde_3775), which is the redox partner of the major pyruvate:ferredoxin oxidoreductase Dde_3237 (Ogata et al., 1988); and most subunits of ATP synthase (although we have insertions in atpA and atpD, encoding the alpha and beta subunits, respectively).

Some genes that are likely to be important for energy production and that we lack data for are Type 1 cytochrome c_3_ (cycA), the soluble cytoplasmic [FeFe] hydrogenase (Dde_0725), and the decarboxylating malate:NADPH dehydrogenase (Dde_1253). Type 1 cytochrome c_3_ is not essential for growth of *D. alaskensis* G20 on lactate/sulfate media but is required for the utilization of hydrogen (Li et al., 2009). The cytoplasmic [FeFe] hydrogenase could be involved in hydrogen cycling or in the utilization of hydrogen as an electron donor.

There are other genes that are absent from the fitness data and are likely to be important for energy production, but in these cases we have fitness data from other genes that are in the same operon and that are expected to be part of the same complex.

## Text S2 — Variable fitness for Hmc and Tmc

Except for the fitness defect for Tmc in hydrogen/sulfate conditions, the fitness values for Hmc and Tmc varied between similar experiments. For example, during formate oxidation, Tmc was important for fitness in two of three experiments (mean fitness −1.0 and −0.59) and Hmc was important in the other one (mean fitness −0.90). The phenotype for Tmc is expected under our model; the phenotype for Hmc is not, but we speculate that it reflects oxidation of hydrogen in the cytoplasm.

The fitness values for Hmc and Tmc were quite variable across 23 lactate/sulfate fitness experiments (mean fitness −0.51 to +0.51 for Hmc and −1.15 to −0.28 for Tmc). However, within each complex, the fitness values were consistent: for example, the lowest pairwise correlation was 0.79 between Hmc genes or 0.90 between Tmc genes. Thus we believe that these phenotypes are genuine.

These experiments were conducted with a variety of conditions, and the variation is partly explained by which base media was used, but we do not know which component of the media was responsible. We used two different base media (“Hazen” or “MO”), at varying concentrations of lactate and sulfate, with or without yeast extract, and with or without added vitamins. Most of the experiments were conducted with 30 mM D,L-lactate, while a few experiments were done with 10 mM lactate. Neither Hmc nor Tmc had much phenotype in the 10 mM lactate experiments, so we removed those experiments before we tried to understand the source of the variation. Among the remaining experiments, 8 were conducted with Hazen media and 50 mM sulfate, and 12 with MO media and 30 mM sulfate. Within each of these groups, yeast extract was added to the media for roughly half of the experiments. Vitamins were added to the media for all of the Hazen media experiments, for all but one of the MO experiments with added yeast extract, and for one of the MO experiments without added yeast extract. The presence of yeast extract did not have a strong effect on the fitness of Hmc or Tmc (both *P >* 0.1, *t* tests). Hmc was slightly detrimental to fitness on MO (mean fitness +0.21, *P* < 0.02, *t* test) while Tmc was important for fitness on MO (mean fitness −0.38, *P* < 0.01). In contrast, on Hazen media, Tmc was very important for fitness (mean fitness −1.2, *P* < 0.01), while Hmc was moderately important (mean fitness −0.35, *P* < 0.01). Among the MO experiments with atypical vitamin usage, we did not see a strong fitness effect of either Hmc or Tmc. Overall, it seems that the base media is a major factor in the variation in fitness.

The major differences between Hazen and MO media were: Hazen media included a higher concentration of sulfate (50 mM versus 30 mM); Hazen media included resazurin (16 nM) as a redox indicator; Hazen media usually included titanium citrate (0.38 mM) as a reductant, while MO media usually included sodium sulfide (1 mM) as a reductant; Hazen media included nitriloacetic acid (0.625 mM) as a chelator while MO media included EDTA (0.12 mM) as a chelator (to reduce precipitation of metal sulfides); Hazen media includes trace minerals at roughly twice the concentration of MO media; and Hazen media was made with a PIPES buffer while MO media was made with a Tris-HCl buffer. In either media, there is sufficient sulfate to reduce 60 mM lactate, but sulfate could become limiting towards the end of growth in MO media. We do not see why the change in buffer or chelating agent or altering the amount of trace minerals would affect the fitness of *hmc* or *tmc* strains. The resazurin and titanium citrate in Hazen media have been suggested to be a stress for Desulfovibrios (J. D. Wall, personal communication), so this might be a relevant difference.

## Acknowledgements

This work conducted by ENIGMA was supported by the Office of Science, Office of Biological and Environmental Research, of the U. S. Department of Energy under Contract No. DE-AC02-05CH11231. The funders had no role in study design, data collection and analysis, decision to publish, or preparation of the manuscript.

We thank Michelle Nguyen, Raquel Tamse, and Ronald W. Davis for microarray hybridization; Yumi Suh and Jil Geller for technical assistance with anaerobic chambers; and Romy Chakraborty and John Coates for sharing equipment.

